# Single-molecule in situ detection and quantification of mitochondrial DNA heteroplasmy

**DOI:** 10.64898/2026.11.18.767531

**Authors:** Sara Gottardo, Suvagata R. Chowdhury, Bryce A. Wilson, Rajini Chandrasegaram, Julien Prudent, Jelle van den Ameele

## Abstract

The mitochondrial DNA (mtDNA) is a multi-copy genome present in most eukaryotic cells and essential for life. The presence of different mtDNA variants within a cell, or heteroplasmy, may cause pathology when the level of deleterious variants exceeds a critical threshold. However, the mechanisms that control heteroplasmy levels remain poorly understood, in part due to a lack of technologies to visualise mtDNA heteroplasmy *in situ* and correlate it with intracellular processes. mtDNA single-molecule fluorescence *in situ* hybridisation (mtDNA-smFISH) allows *in situ* visualisation of mtDNA molecules and sequence-specific detection of heteroplasmic variants. Here, we provide protocols for sequence-specific detection of mtDNA, large-scale single mtDNA deletions and species-specific haplotypes, in fixed human and *Drosophila* cells and in whole-mount *Drosophila* tissues. Combining mtDNA-smFISH with immunohistochemistry using biotin-labelled primary antibodies that resist heat-based denaturation, allows concomitant detection of intracellular proteins. We further describe two image-analysis pipelines, one for 3D quantification and one open-source for 2D quantification of mtDNA deletions in *Drosophila* tissues and cell culture. This enables estimation of single-cell heteroplasmy levels, observation of cell-to-cell variability, and detection of mtDNA variants at a subcellular level, making mtDNA-smFISH a powerful tool to study the intracellular processes regulating heteroplasmy and mtDNA quality control.

## 1 Introduction

Mitochondria are critical double-membrane organelles, involved in the generation of cellular energy and a broad range of cellular homeostatic processes, as well as in the control of cell fate decisions. ^1^ Unlike all other organelles in animal cells, mitochondria are unique, since they contain their own genome. Mitochondrial DNA (mtDNA) consists of a double-stranded circular molecule of about 16.5 kb in humans, containing 37 genes: 13 encoding for OXPHOS complex subunits, 22 for tRNAs, and 2 for rRNA. ^2^ In contrast to the two copies of nuclear DNA, mtDNA exists in multiple copies in a single cell, with copy number ranging from hundreds to thousands of molecules depending on cell type, and replicates independently of the cell cyle. When a mutation arises in mtDNA, it usually affects only a fraction of all molecules within a cell. The coexistence of wild-type (WT) and mutated mtDNA within the same cell is referred to as heteroplasmy. ^3,4^ Levels of heteroplasmy vary greatly between cells within a tissue, across different tissues and organs in the same organism, and between organisms across generations. ^5^ While low levels of heteroplasmy are found in all human tissues and cells, mutation load can also shift throughout an individual’s life. Ageing and age-associated pathologies, including cancer and neurodegenerative diseases, have all been linked to this accumulation of mtDNA mutations within cells over time. ^6–9^

Variability in heteroplasmy is also associated with primary mitochondrial diseases, a group of genetic disorders characterised by highly heterogeneous clinical manifestations. ^10,11^ When the mutation load in specific cells or tissues reaches pathogenic thresholds, it triggers mitochondrial dysfunction and symptom manifestations that vary substantially between patients. ^4,12^ Studies focusing on the processes regulating variability and shifts in heteroplasmy encounter major technical challenges. While single-cell sequencing and genotyping approaches provide valuable insights, they require tissue dissociation and analysis of isolated cells. ^8,13,14^ In doing so, key information about the distribution of a mutation and its correlation with the tissue architecture are lost. However, spatial information, within a tissue and at the subcellular level, is likely to provide essential insight into the complex mechanisms driving and regulating mtDNA heteroplasmy dynamics, ageing and disease. A range of techniques has previously been proposed for sequence-specific imaging of mtDNA and detection of heteroplasmic variants. ^14–25^ However, each has its own advantages and limitations. Standard Fluorescence *In Situ* Hybridization (FISH) has previously been used successfully for labelling and imaging large-scale mtDNA variation in cell culture ^25^ and whole-mount *Drosophila* tissues ^16^, but relies on large probes with limited sensitivity and resolution to distinguish single mtDNA molecules. More versatile approaches for *in situ* visualisation of heteroplasmic mtDNA variants, including in somatic tissues, are therefore needed to understand the cellular and molecular processes that cause somatic heteroplasmy variation and tissue-specific phenotypes of mitochondrial disease, and to answer long-standing questions in the field of mitochondrial and mtDNA biology. ^15^

Here, we describe an effective strategy for sequence-specific microscopic visualisation of mtDNA in human and *Drosophila* fixed cells , and in whole-mount *Drosophila* tissue. Our approach builds on single-molecule Fluorescent *in situ* Hybridisation (smFISH) in combination with Hybridisation Chain Reaction (HCR). ^26,27–28^ smFISH-HCR has been implemented widely as a powerful and versatile approach for straightforward, sensitive and specific *in situ* detection of RNA molecules in a range of experimental contexts. We adapted smFISH-HCR for *in situ* detection of mtDNA, ^29^ and here outline protocols to apply this in fixed cultured human or *Drosophila* cells and whole-mount *Drosophila* tissue, and to combine mtDNA-smFISH with immunostaining, including against antigens destroyed by the harsh denaturation conditions. We further provide probe sequences to distinguish *D. melanogaster* and *D. Yakuba* mtDNA variants in heteroplasmic *Drosophila* strains carrying a combination of both mtDNA genotypes within the same cells; and probes to distinguish full-length mtDNA molecules from those carrying a large-scale deletion in cultured human cybrid cells. This protocol performs well in the presence of substantial sequence differences, however, it has not yet been optimised for detection of single-nucleotide polymorphisms. Finally, we outline approaches for 3D automated or manual heteroplasmy quantification, and describe an open-access 2D image analysis pipeline for automated smFISH-analysis and estimation of heteroplasmy levels.

mtDNA-smFISH has proven useful to investigate mtDNA dynamics in *Drosophila*, confirming the presence of purifying selection against pathogenic mtDNA variants during oocyte development and identifying presence of a somatic bottleneck accompanied by purifying selection during neurogenesis in the developing *Drosophila* brain. ^29^ Going forward, this strategy provides a powerful platform to investigate the spatial distribution, cell-to-cell variability, and tissue specific dynamics of mitochondrial heteroplasmy, while also offering new opportunities to analyse how heteroplasmic mtDNA molecules behave during different cellular processes and mtDNA behaviours, including mtDNA degradation, trafficking and release. Together, these applications will deepen our knowledge of mitochondrial genetics and mtDNA cell biology in development, ageing and disease.

## 2 Materials (*see* Note 1)

### 2.1 Common disposables

- 12-well chamber, removable (Ibidi, #81201)
- Tissue culture dish 100 mm, Standard (Sarstedt, #83.3902)

### 2.2 Experimental models

#### 2.2.1 HeLa cells

Obtained from American Type Culture Collection (ATCC CCL-2).

#### 2.2.2 ΔH2.1 cybrid cell line

Obtained from Carlos Moraes, carrying heteroplasmic 7.5 kb mtDNA deletion from m.7982 to m.15504. ^30^

#### 2.2.3 Drosophila melanogaster cells

*Drosophila* Genomics Resource Centre stock number 150, FlyBase ID FBtc0000150.

#### 2.2.4 Drosophila melanogaster strains

All maintained at 18-29 °C, unless specified otherwise.

- Homoplasmic *D. melanogaster*, *w*^1118^*;+;+*. Bloomington Drosophila Stock Centre (BDSC) stock number 5905
- Heteroplasmic *Drosophila*, carrying a combination of WT *D. yakuba* mtDNA and mutant *D. melanogaster* mtDNA (mt:ND2del1+mt:CoIT300I), on a nuclear *D. melanogaster* w^1118^ genomic background, as described previously. ^31^ Obtained from Hansong Ma. This strain has to be maintained at 29 °C, in order to ensure stable heteroplasmy levels. Heteroplasmy can be monitored routinely by PCR-restriction digest, as described previously. ^29^
- Homoplasmic *Drosophila* carrying *D. yakuba* mtDNA on a nuclear *D. melanogaster* w^1118^ genomic background, as described previously. ^32^ Obtained from Hansong Ma.
- UAS-Mito-XhoI-Myc transgenic strain for expression of a mitochondrial-targeted restriction enzyme. ^33^ BDSC stock number 81007
- *worniu*-GAL4 transgenic strain on chromosome II, for neural stem cell (NSC)-specific transgene expression. ^34^ BDSC stock number 56553

### 2.3 *Drosophila* husbandry

- Standard cornmeal agar medium supplemented with dry yeast

### 2.4 Cell Culture

- Dulbecco’s Modified Eagle Medium (Thermo Fisher Scientific, #10566016)
- Schneider’s *Drosophila* Medium (Thermo Fisher Scientific 21720024)
- Fetal Bovine Serum (FBS) (Thermo Fisher Scientific, #16140071)
- Trypsin (Sigma-Aldrich, #T2601)
- Uridine (Sigma-Aldrich, #U3003)

### 2.5 mtDNA-smFISH probes and reagents

- Formaldehyde, Methanol-free (Thermo Fisher Scientific, #28908)
- Paraformaldehyde (Sigma-Aldrich, #P6148)
- Formamide Deionized (Merck Millipore, #S4117)
- TritonX-100 (Sigma-Aldrich, #STBJ7635)
- Tween20 (Sigma-Aldrich, #SLCH9711)
- 20X saline-sodium citrate (SSC) buffer (Sigma-Aldrich, #S6639)
- Probe hybridisation buffer (Molecular Instruments, #BPH02023) (*see* **Note 2**)
- Probe wash buffer (Molecular Instruments, #BPW002225) (*see* **Note 2**)
- Amplification buffer (Molecular Instruments, #BAM02622) (*see* **Note 2**)
- Amplifier hairpins (Molecular Instruments)
- Probes (Molecular Instruments, *see* **Section 3.1** and **Table 1**)

### 2.6 Immunofluorescence reagents and antibodies

- mouse anti-ATPsynα (Abcam #ab14748)
- rabbit anti-FLAG M2 (Cell Signalling #D6W5B)
- chicken anti-GFP (Abcam **#**ab13970)
- guinea pig anti-heat denatured GFP (Kouichi C. Nakamura and Hiroyuki Hioki) ^35^
- rabbit anti-heat denatured GFP (Kouichi C. Nakamura and Hiroyuki Hioki) ^35^
- rabbit anti-TFAM (GeneTex #GTX103231)
- rabbit anti-TOMM20 (Thermo Fisher Scientific #10G6I8)
- mouse Ig2a anti-TOMM20 (Santa Cruz #sc17764)
- biotin-conjugated goat anti-Chicken IgY (H+L) secondary antibody (Invitrogen #PA1-28657)
- Alexa Fluor 488 Streptavidin (from Abcam Streptavidin AZDye™ kit #ab272187) or conjugated to other fluorophores
- Alexa Fluor goat anti-rabbit IgG (H + L) Alexa Fluor 647 (Thermo Fisher Scientific #A-21244)
- Alexa Fluor Goat anti-mouse IgG cross-Adsorbed Alexa Fluor 647 (Thermo Fisher Scientific #A-21235)
- Alexa Fluor Goat anti-rabbit IgG2a cross-Adsorbed Alexa Fluor 647 (Thermo Fisher Scientific #A-21241)
- Prolong Diamond antifade mounting medium (Thermo Fisher Scientific #P36961)
- DAKO Fluorescence Mounting medium (Agilent Technologies #S3023)

### 2.7 Other reagents

- 2’-3’-dideoxycytidine (Sigma-Aldrich #D5782)
- RNAse A (Roche #10109142001)
- Bovine Albumine Fraction (BSA) (Merck #A6003)
- Nde I (NEB #R0111)

### 2.8 Equipment

- Thermoblock (Labnet)
- Incubator for human cell culture, maintained at 5% CO2 and 37 °C
- Incubator for *Drosophila* cell culture, humidified and maintained at 25 °C
- Incubator for *Drosophila* animals, to maintain strains at 18 °C, 25 °C and 29 °C
- Zeiss LSM880 confocal microscope with Fast AiryScan or equivalent
- Nikon Eclipse TiE with Andor Dragonfly 500 spinning disk system and Zyla 4.2 PLUS sCMOS camera or equivalent

### 2.9 Software

- Fiji (ImageJ, v1.53f or later)
- MorphoLibJ
- FishQuant

## 3 Methods

The protocol steps detailed here apply to both cell culture and whole-mount tissue samples. In the few instances where model-specific considerations apply, these are explicity stated in the relevant step. Steps 3.3.3 and 3.4, provide instructions for combining immunostaining with mtDNA-smFISH, are optional and can easily be skipped otherwise, without further modifications to the protocol.

**Table 1:**
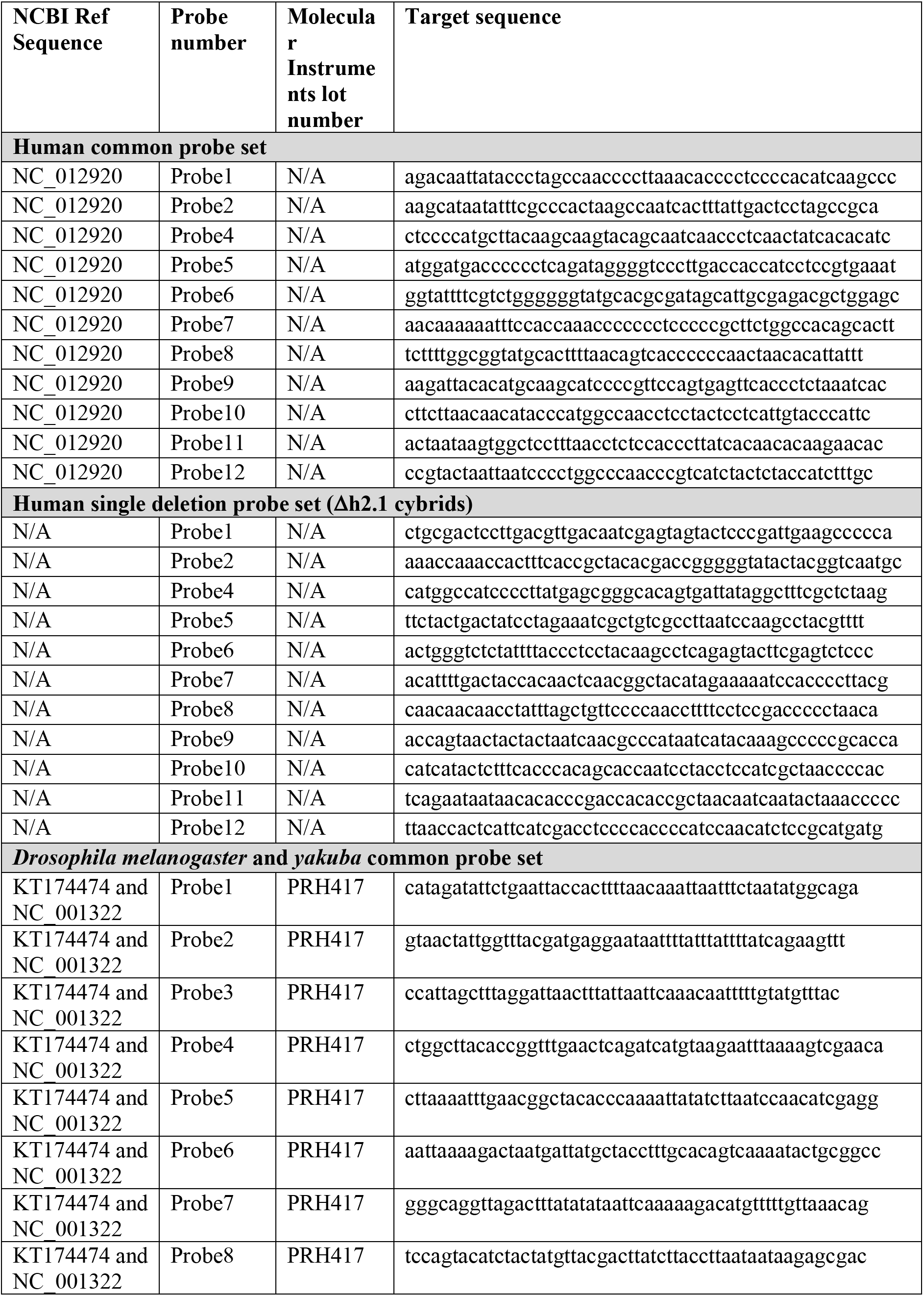
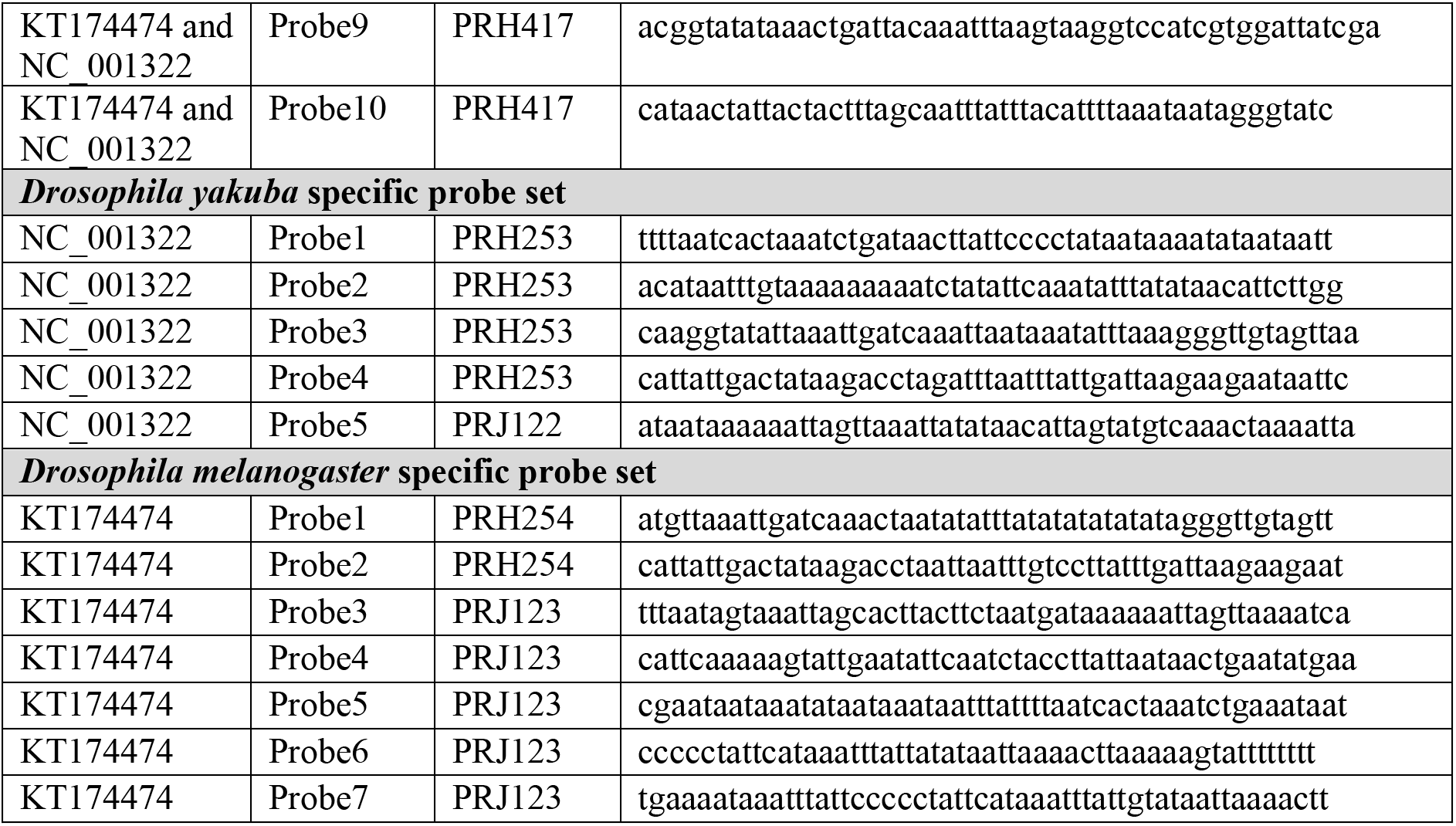
smFISH-mtDNA probe target sequences. All sequences provided are 5’-3’.

### 3.1 Probe design

1. mtDNA-smFISH is based on smFISH-HCR v3.0. ^26,27^ It relies on pairs of short single-stranded DNA probes, whereby each probe half recognises a 25-nucleotide target sequence, with two nucleotides spacing between both sides of a pair (**Figure 1A**). The probes further contain a short two-nucleotide spacer sequence, and an 18-nucleotide overhang. The overhang of each probe provides half of the initiator sequence for the HCR reaction, which is only triggered when both halves of the initiator sequence are positioned in close proximity, through specific binding of the target mtDNA sequence (**Figure 1B**).
2. Probes can be ordered from a commercial provider (*see* Reagents and **Note 3**), can be designed manually or generated using an open-source HCR probe-design software. ^36^ When designing probes manually, the probe sequence is the reverse complement of the target sequence (*see* **Note 4**). Each probe half can then be designed together with its initiator sequence (I1 or I2), which is determined by the choice of HCR amplifier sequences. The probe sequence is assembled as follows (*see* **Note 5** for description of initiator sequences):
3. Probe half 1: <u>Initiator1</u>-**Initiator1spacer**-Reversed5’sideProbe
4. Probe half 2: Reversed3’sideProbe-**Initiator2spacer**-*Initiator2*
5. Probe design for an example target sequence
  - 52-nucleotide target sequence (with the 2-nucleotide spacer sequence in bold): 5’-TACAACTACAACAGCCACAACGTCT**AT**ATCATGGCCGACAAGCAGAAG AACG-3’
  - Initiator sequence, for example for the B1 HCR amplifier sequence, with the 5’-half for Probe 1 underlined, and the 3’-half for Probe 2 in italics: 5’-GAGGAGGGCAGCAAACGG*GAAGAGTCTTCCTTTACG*-3’
  - Probe 1: 5’-GAGGAGGGCAGCAAACGG**AA**AGACGTTGTGGCTGTTGTAGTTGTA-3’
  - Probe 2: 5’-CGTTCTTCTGCTTGTCGGCCATGAT**TA***GAAGAGTCTTCCTTTACG*-3’
6. 11 and 10 probe pairs were designed to detect human and *Drosophila* mtDNA, respectively (**Table 1**). Probes can be designed to assess heteroplasmy by targeting regions of the mtDNA that differ sufficiently between wild type and mutant DNA (*see* **Notes 6, 7** for a discussion of choosing target sequences).
7. In the case of ΔH2.1 cybrid cell line, a set of 11 probe pairs was designed to target sequences contained in the 7.5 kb deleted sequence (**Figure 1C**; **Table 1**). This “deletion probe set” will only recognise full-length molecules of mtDNA. Differential visualisation of the mutant vs. WT mtDNA then relies on the overlap of the signal from the deletion and common probe sets, whereby the WT mtDNA is labelled by both, and the deleted only by the common probe set (**Figure 1D**).
8. To distinguish *D. melanogaster* and *D. yakuba* mtDNA molecules, sets of 7 and 5 probes were designed respectively (*see* **Note 7**; **Table 1**).
9. It is important to consider conditions to test probe specificity for the experimental system of choice. For a discussion of testing specificity using DNase or RNase treatment, *see* **Note 8**; for other conditions in cell culture, *see* **Note 9**; and in whole-mount *Drosophila* tissue, *see* **Notes 10, 11**.

### 3.2 Experimental models

#### 3.2.1 Human cell culture

Cells used are HeLa cells or ΔH2.1 cybrid cells ^30^. Culture cells in Dulbecco’s Modified Eagle Medium (DMEM) supplemented with 10% FBS at 5% CO_2_ and 37 °C. When working with ΔH2.1 cybrid cell lines, add 50 µg/mL uridine to the medium.

**Figure 1.**
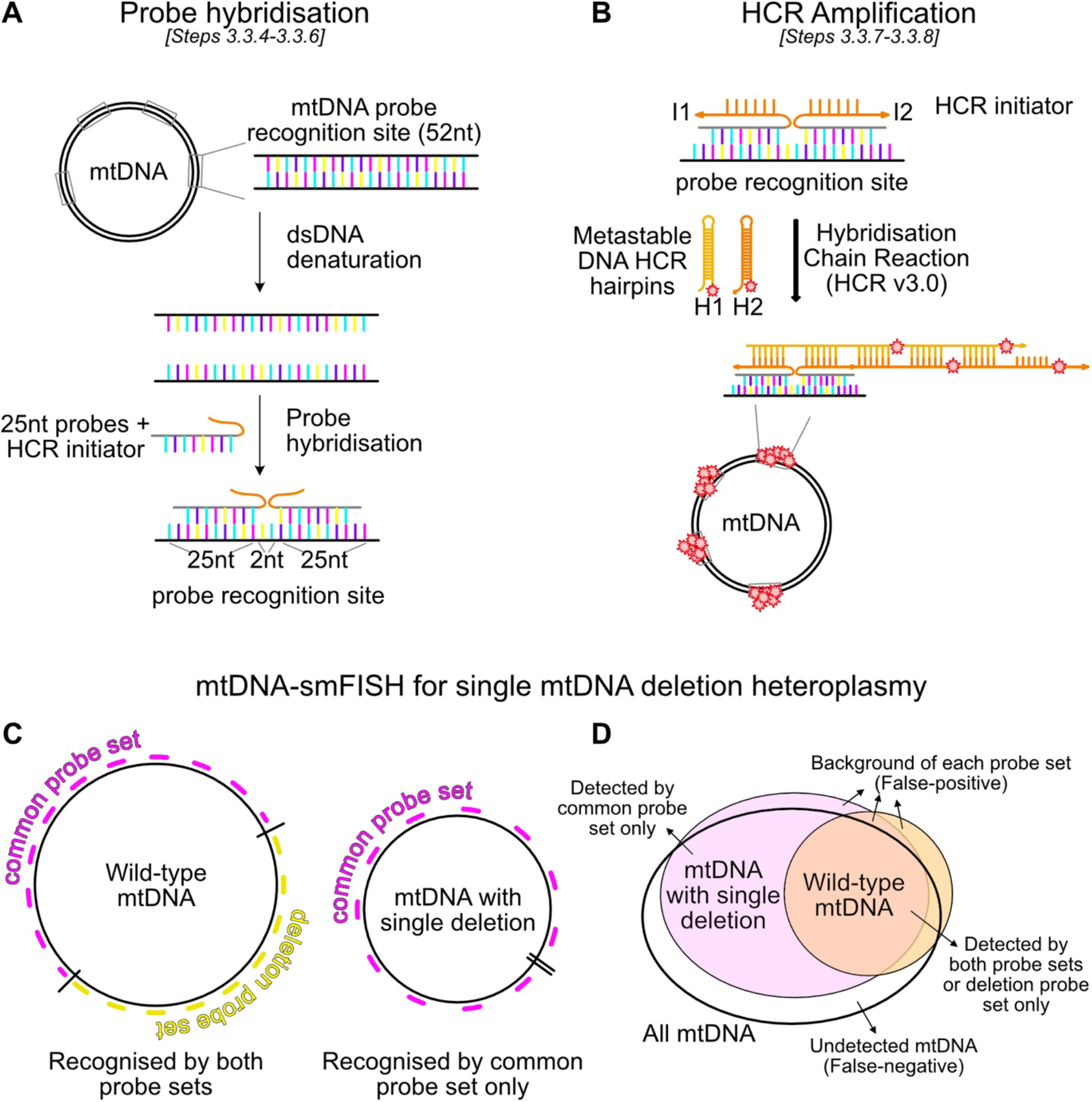
mtDNA-smFISH protocol. **A.** Schematic of mtDNA-smFISH probe design, as two pairs of 25-nucleotide sequences separated by 2 nucleotides, together targeting 52-nt long mtDNA sequences, as detailed in **Section 3.1**. *In situ* probe hybridisation (Section 3.3.5) occurs after denaturation (Section 3.3.4) of the double-stranded target mtDNA, as detailed in **Section 3.3**. **B.** During hybridisation chain reaction (HCR), fluorescent hairpins bind to the initiator sequence present on the probes, and signal is amplified as the hairpin chain elongates, as detailed in **Section 3.3.7-3.3.8**. **C-D.** Schematic of probes designed against sequences within (deletion probe set) and outside (common probe set) a human 7.5kb single mtDNA deletion (C), allowing differential visualisation of WT and mutant mtDNA molecules with mtDNA-smFISH (D), as detailed in **Section 3.1** and **3.7**. Schematics adapted, with permission, from Ref. ^29^, under a Creative Commons Attribution 4.0 International License.

#### 3.2.2 *Drosophila* cell culture

Culture cells in Schneider’s *Drosophila* Medium supplemented with 10% FBS, in a humidified incubator at 25 °C. When cells reach 80-90% confluency, they are scraped and mixed, and 25% of the cell suspension is transferred into fresh culture flasks.

#### 3.2.3 *Drosophila* husbandry

Homoplasmic *Drosophila* stocks can be maintained at 18 or 25 °C on standard cornmeal food in plastic vials with dried yeast. Heteroplasmic *Drosophila* lines need to be maintained at 29 °C. The presence of heteroplasmy can be validated by PCR and restriction digest (*see* **Note 12**).

### 3.3 mtDNA-smFISH

#### 3.3.1 Sample preparation and Fixation

• Human and *Drosophila* cell culture: wash cells in pre-warmed PBS and detach by adding trypsin (human cells) or by scraping (*Drosophila* cells). Seed about 20,000 cells per well of a 12-well chamber (removable) (*see* **Note 13**). The maximum volume for each well is 250 µL, so 200 µL is the volume used for each solution in the following steps, unless otherwise specified (*see* **Note 14**). After 24 hours, remove the medium and fix the cells by adding 5% paraformaldehyde (powder dissolved in PBS) at 4 °C for 20 minutes. Wash three times in PBS with 0.1% Tween-20 for 5 minutes. Representative images of mtDNA-smFISH in cultured cells are provided in **Figure 2**.

**Figure 2.**
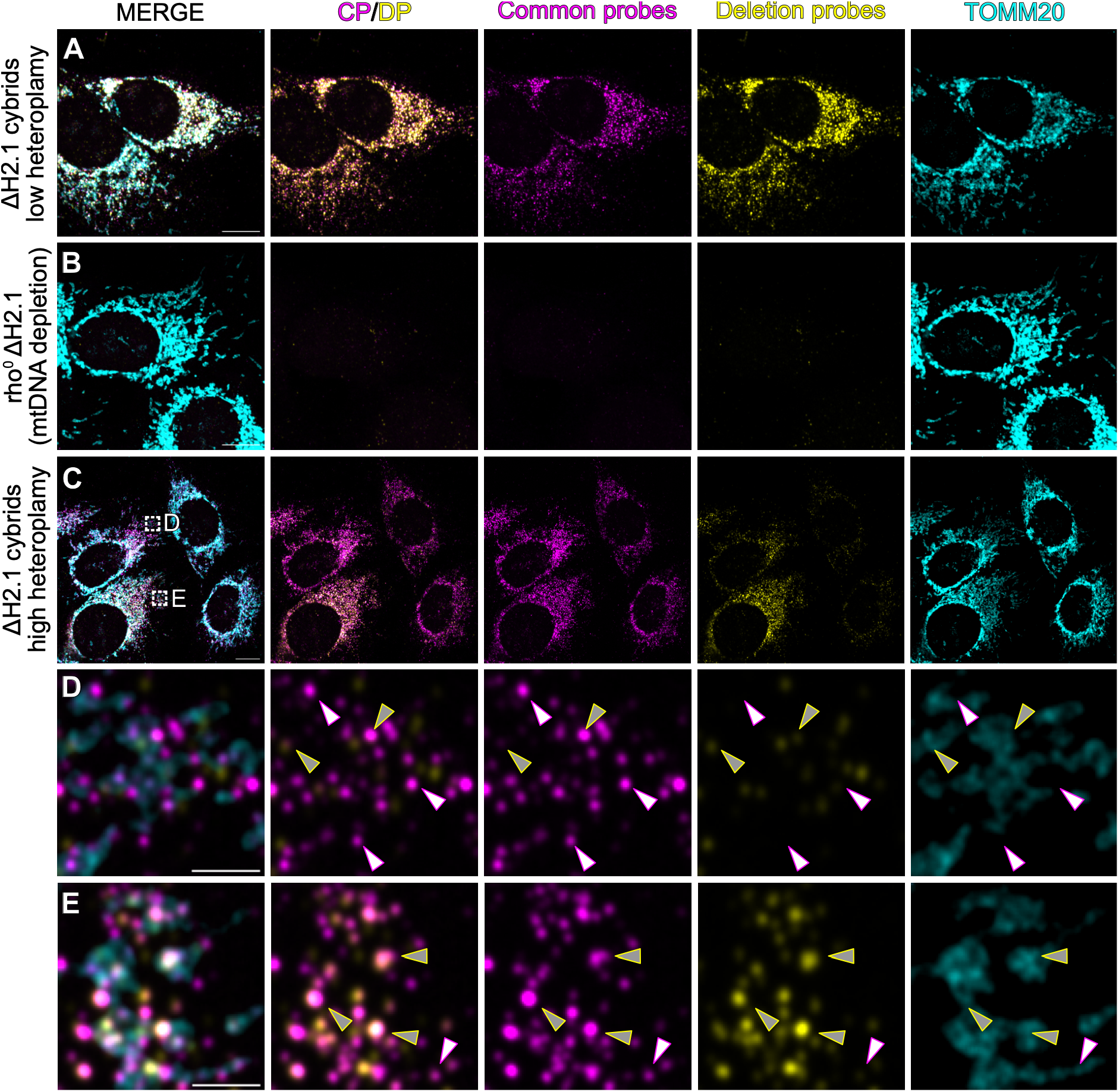
mtDNA-smFISH in cultured cells A-E. mtDNA-smFISH on ΔH2.1 cybrid cells with common and deletion probe sets (*see* Figure 1C**,D**), along with standard immunostaining for TOMM20, as detailed in **Section 3**. ΔH2.1 cybrid cells with low mutant heteroplasmy (**A)** have much higher deletion probe (DP) signal relative to common probe (CP) signal. Both CP and DP signal disappear upon mtDNA depletion and the generation of Rho^0^ cybrid cells (**B**) (*see* **Note 9**). Within high heteroplasmy cells (**C-E**), variability in heteroplasmy levels can be seen, with adjacent cells having more (**D**) or less (**E**) CP puncta (magenta) without DP (yellow) overlap, indicating relatively lower heteroplasmy levels in (E) (*see* Figure 1D for a schematic). White arrowheads, CP-only puncta; grey arrowheads, CP/DP puncta. Scale bars are 10μm (A) or 2μm (D,E).

• Whole-mount *Drosophila* tissue: perform tissue dissections (*see* **Note 15**) in PBS and fix in 4% formaldehyde (diluted 1/4 from 16% stock-solution) in PBS with 0.3% Triton X-100 for 30 minutes at room temperature. Wash three times in PBS with 0.1% Tween-20 for 5 minutes. Representative images of mtDNA-smFISH in whole-mount *Drosophila* tissue are provided in **Figure 3**.

**Figure 3.**
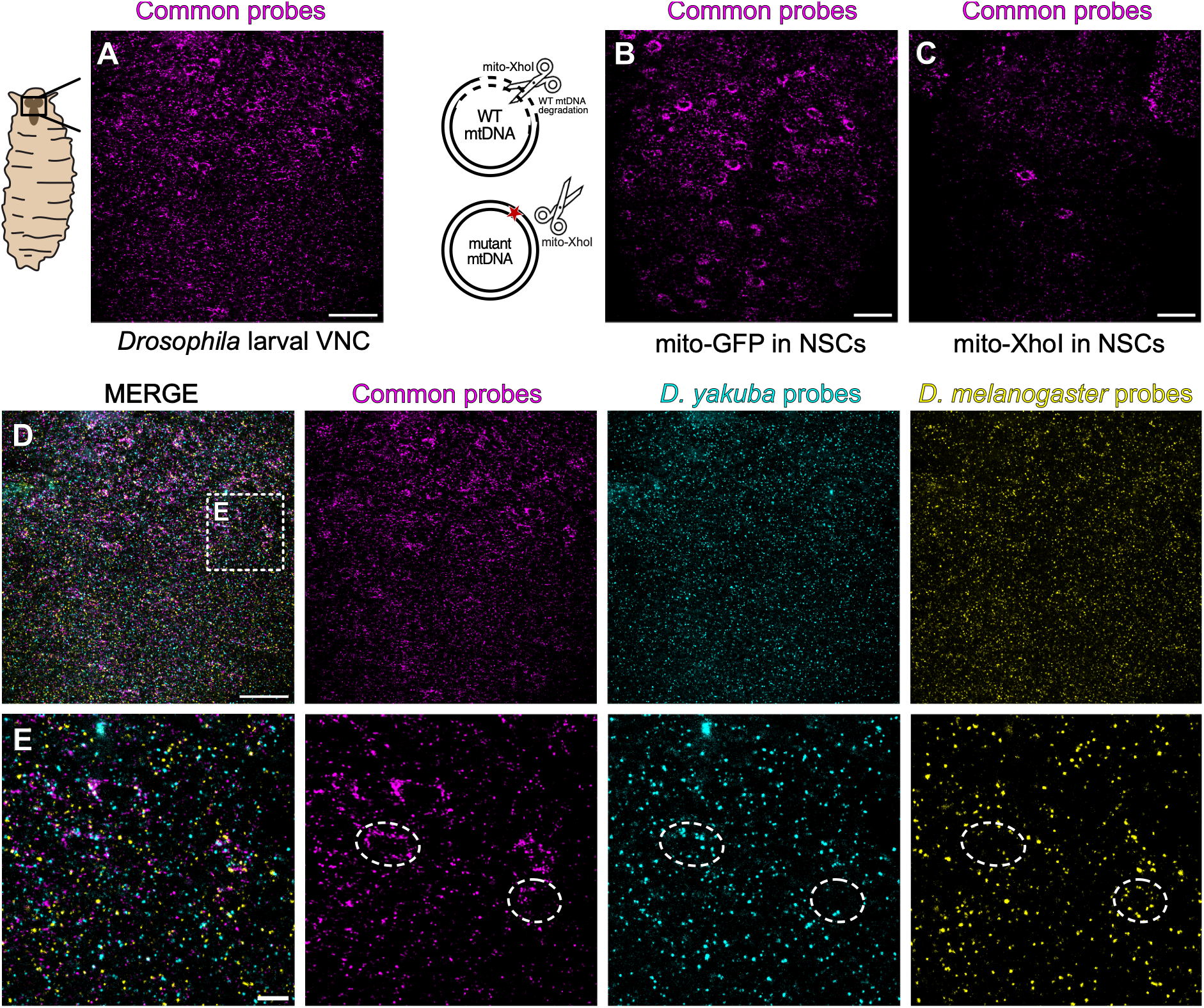
mtDNA-smFISH in whole-mount *Drosophila* tissue. **A.** mtDNA-smFISH on whole-mount *Drosophila* larval brain with common probe set, as detailed in **Sections 3.1** and **3.3**. **B-C.** Depletion of common probe mtDNA signal in *Drosophila* brains following NSC-specific (worniu-Gal4) expression of mitochondrial targeted XhoI **(C)** with mitochondrial targeted GFP **(B)** as a control. WT *Drosophila* mtDNA is degraded following cleavage by XhoI (*see* **Note 10**). **D-E.** mtDNA-smFISH in heteroplasmic *Drosophila* larval brain with *D. yakuba* (WT, cyan), *D. melanogaster* (mutant, yellow), and common probe (magenta) sets. Variability in heteroplasmy levels can be seen in adjacent NSCs (E). Scale bars are 25μm (A-D) and 5μm (E). Schematics reproduced, with permission, from Ref. ^29^, under a Creative Commons Attribution 4.0 International License.

#### 3.3.2 De/rehydration

1. Dehydrate the sample by performing serial 5-minute immersions in ice cold (4 °C, on ice) methanol (MeOH) in PBS with 0.1% Tween-20 with increasing MeOH concentrations: first 25% MeOH in PBS with 0.1% Tween-20, then 50% MeOH, 75% MeOH and finally 100% MeOH. Once samples are in 100% MeOH, they can be stored at -20 °C.
2. Follow these steps in reverse order (from 100% MeOH to 0% MeOH in PBS with 0.1% Tween-20) to rehydrate the sample, shortly before proceeding to the next mtDNA-smFISH steps.

#### 3.3.3 (Optional) Immunostaining with antigen biotinylation

If immunostaining is required against an antigen that does not withstand heat denaturation (*see* **Note 16**), additional steps are performed before proceeding to denaturation (**Figure 4**). The steps below are an example of standard immunostaining for cells or whole-mount tissues, but can be adapted to any suitable immunostaining protocol. For immunostaining against antigens that can withstand heat denaturation (*see* **Note 17**), these steps should *not* be performed (*see* **Section 3.4**).

1. After de/rehydration (*see* **Section 3.3.2**), perform three 5-minute washes in PBS/Triton X-100 0.1%.
2. (Optional) A blocking step can be performed by incubating samples in PBS/Triton X-100 0.1% with 10% FBS and/or 3% BSA for 20 minutes at room temperature. This depends on the experimental conditions and primary antibody.
3. Incubate samples overnight at 4 °C with the primary antibody of choice in PBS/Triton X-100 0.1%.
4. Wash samples three times for 10 minutes in PBS/Triton X-100 0.1% .
5. Incubate samples overnight at 4 °C with a biotin-conjugated secondary antibody in PBS/Triton 0.1%.
6. Wash samples three times for 10 minutes in PBS/Triton 0.1%.
7. Perform another fixation step with 4% FA in PBS/Triton X-100 0.1% for 20 minutes at room temperature. This second fixation is required to ensure that the antibodies remain in place during the subsequent harsh denaturation conditions and the subsequent washes.
8. Wash samples three times for 10 minutes in PBS/Triton X-100 0.1%.

**Figure 4.**
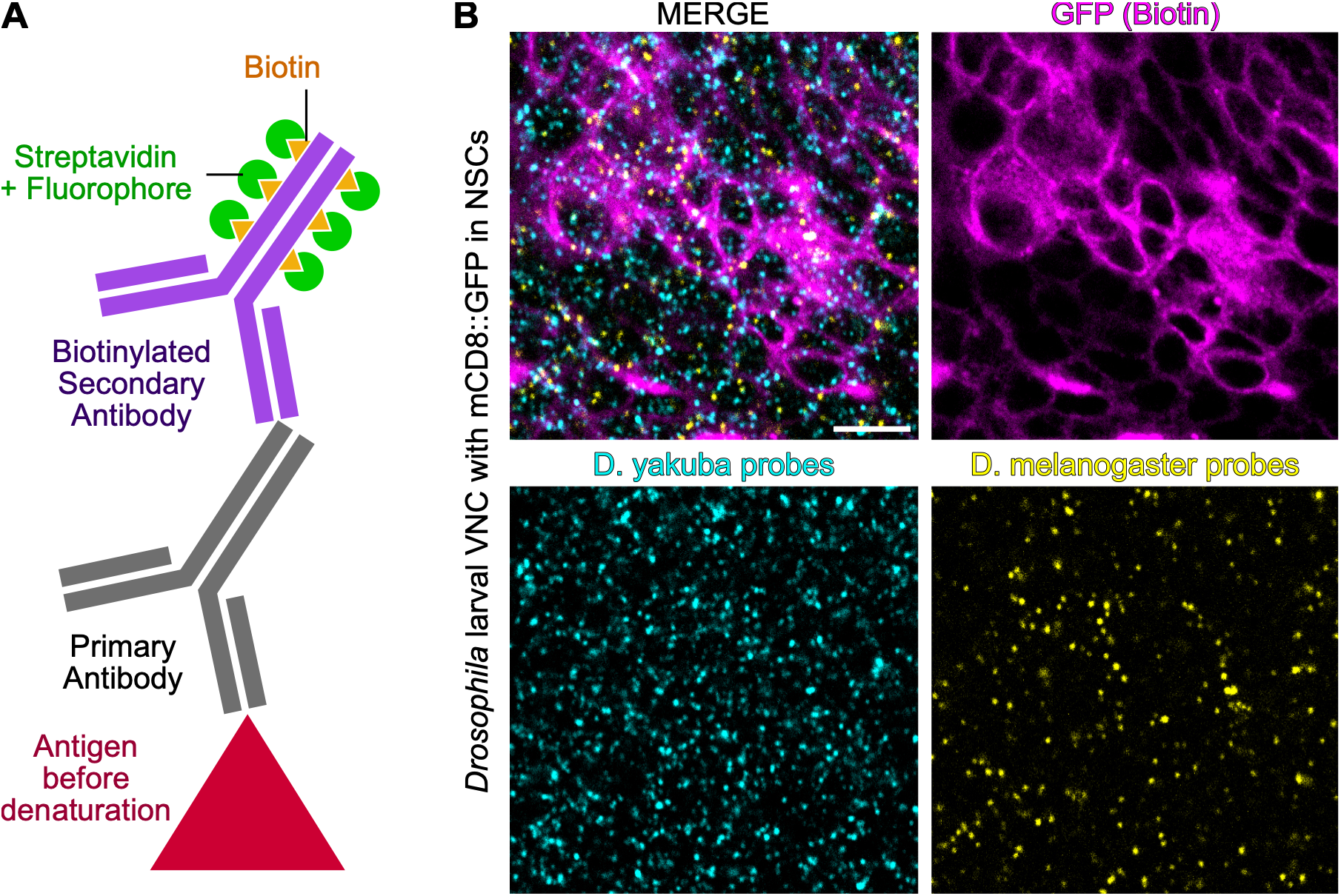
Immunostaining with biotinylated secondary antibodies. **A**. Prior to the mtDNA-smFISH denaturation step, normal primary antibody staining can be performed along with a biotinylated secondary antibody, as detailed in **Sections 3.3.3** and **3.4**, and **Note 16**. **B**. Representative images of biotinylated immunostaining against GFP performed with mtDNA-smFISH on a heteroplasmic *Drosophila* larval brain expressing membrane targeted GFP (mCD8::GFP) in NSCs (*Worniu-GAL4*) and their progeny. Scale bar is 5μm.

#### 3.3.4 Denaturation

1. Prepare denaturation solution with 70% formamide and 2x SSC in ultra-pure water.
2. Remove PBS with 0.1% Tween-20 solution from the last rehydration step and add denaturation solution.
3. Heat samples in denaturation solution at 90 °C for 10 minutes.
  - For cells, using a thermoblock with the heating block turned upside-down is recommended, with the chamber placed on the flat surface of the block, and covered by a polystyrene box.
  - For whole-mount tissues, vials containing tissues in denaturation solution are placed in a shaking heating block at 300 rpm.

#### 3.3.5 Hybridisation

1. Dilute probes at 4 nM in hybridisation buffer. 4 nM is the concentration of each individual probe pair, not the total concentration of all probes combined. For lower probe numbers, higher concentrations of probes can be used, but this should be determined empirically based on the signal intensity obtained at the end of the protocol. Typically, we increase the probe concentration to 20 nM for fewer than 10 probes, and have obtained acceptable results at 100 nM for single probe pairs (*see* **Note 18**).
2. Heat probes in hybridisation buffer to 90 °C for 2 minutes
3. Carefully remove the denaturation solution from the cells or tissues, making sure not to aspirate tissues that may float or stick to the side of the vials. Minimise the amount of time each vial is kept outside the heat block, and place the samples back on or in the heat block immediately after removing the denaturation solution.
4. Add the probes in hybridisation buffer, again minimising the amount of time spent outside the heat block
5. Remove from heat and incubate at 37 °C overnight, using a shaking heat block for tissue samples. Alternatively, lower the temperature of the heat block to 37 °C, and let the samples cool in the heat block before leaving them there overnight. For cell culture, keep the 12-well chamber in a humid environment by placing it in a container with wet tissue paper, to avoid evaporation.

#### 3.3.6 Post-hybridisation washes

1. Warm probe wash buffer to 37 °C.
2. Perform four washes of 15 minutes each with pre-warmed probe wash buffer. For whole-mount tissue samples, adding 200 µl of wash buffer first to the sample while it is still in hybridisation buffer will help the sample to sink, making it easier to remove the hybridisation buffer without tissue loss.
3. While performing the washes, defrost the HCR hairpins while keeping them in the dark. Make sure to thaw both hairpins (H1 and H2) for each amplification reaction, and use the appropriate hairpin sequences that match the initiator sequences of the probes.
4. Heat the thermoblock to 95 °C.
5. Warm the required amount of amplification buffer (50 µl per well for cells; 100 µl per sample for whole-mount tissue) to room temperature.

#### 3.3.7 HCR-mediated amplification

1. During the fourth probe wash buffer wash, anneal HCR hairpins by heating and subsequent snap-cooling, to ensure intramolecular folding of each hairpin. Use 1 µL (3 pmol, of a 3 µM stock) per well of each hairpin for cells, or 2 µl (6 pmol, of a 3 µM stock) per sample of each hairpin for whole-mount tissue, and place the required amount of each hairpin separately into a vial. For example, for a full cell culture 12-well chamber, place 12 µl of hairpin 1 in one vial, and 12 µl of hairpin 2 in a second vial. Heat the vials at 95 °C for 90 seconds and snap-cool hairpins by moving them to room temperature for 30 minutes, while keeping them in the dark. Do *not* combine H1 and H2 hairpins at this stage, as this may cause them to anneal to each other.
2. While hairpins cool down, wash samples for 5 minutes with a solution composed of equal parts of probe wash buffer and 5x SSC-PBS/Tween-20 0.1% at room temperature.
3. Continue with two 5 minutes washes in 5x SSC-PBS/Tween-20 0.1% at room temperature.
4. Dilute the hairpins in amplification buffer to a 60 nM concentration: 1 µl of a 3 µM stock per 50 µL of amplification buffer for cells; 2 µl in 100 µl buffer for whole-mount tissue. For example, add 12 µl of hairpin 1 and 12 µl of hairpin 2 to a total of 600 µl for a full 12-well chamber.
5. Remove the last wash and add the premixed hairpins in amplification buffer to the sample.
6. Incubate at room temperature in the dark for 6-24 hours (*see* **Note 19**). For cells, during this incubation, a humid environment helps avoid evaporation.

#### 3.3.8 Post-amplification washing

1. Perform two washes with 5x SSC-PBS/Tween-20 0.1% (SSCT) for 5 minutes each, two washes of 30 minutes, and one final wash of 5 minutes. Continue by performing three 10-minute washes with PBS with 0.1% Triton X-100, and finally two 10-minute washes with PBS. If immunostaining will not be performed, proceed to step 6, Mounting.
2. If immunostaining will be performed after mtDNA-smFISH, perform serial washes to gradually replace 5x SSC-PBS/Tween-20 0.1% (SSCT) with PBS/Triton X-100 0.1%. Start with a first 5-minute wash with 75% SSCT in PBS/Triton X-100 0.1%. For the second wash, reduce SSCT to 50%. In the last wash, use 25% SSCT. Finally, perform three washes of 5 minutes with PBS/Triton X-100 0.1% or a buffer that will be used for immunostaining, and proceed with **Section 3.4**.

#### 3.4 (Optional) Immunostaining

##### 3.4.1 Immunostaining against antigens that withstand heat denaturation (*see* **Notes 16, 17**)

1. Proceed with any suitable immunostaining protocol. The steps below are an example of a standard immunostaining for cells or whole-mount tissues. The same steps can be used for antibodies raised specifically against heat-denatured antigens, such as for GFP. ^35^
2. 2. An optional blocking step can be performed by incubating samples in PBS with 10% FBS and/or 3% BSA for 20 minutes at room temperature.
3. 3. Incubate samples overnight at 4 °C with primary antibodies diluted to the appropriate concentration in PBS with 5% FBS.
4. 4. Wash samples with PBS three times.
5. 5. Incubate samples with secondary antibodies at the appropriate dilution in PBS with 5% FBS for 1 hour at room temperature.

##### 3.4.2 Streptavidin-based detection of biotinylated antibodies (*see* **Note 17**)

The first eight steps of immunostaining are performed prior to denaturation (*see* **Section 3.3.3**), and the immunostaining steps are then finished after the post-amplification washing.

1. After the mtDNA-smFISH post-amplification washing steps are completed, incubate samples overnight at 4 °C with fluorescently labelled streptavidin. Streptavidin will bind to the biotin conjugated to the secondary antibody (*see* **Section 3.3.3**), since biotin was not destroyed during the denaturation steps. ^29,37^
2. Wash samples three times for 10 minutes each in PBS/Triton X-100 0.1%.

### 3.5 Mounting

• Cell slides are washed with PBS three times and once more with deionised water before mounting. Remove the wells of the chamber, apply mounting medium (*see* **Note 20**) and mount the coverslip. Press gently to remove air bubbles.

• Whole-mount tissues are mounted (*see* **Note 20**) according to standard laboratory practices.

### 3.6 Image acquisition

1. For imaging of human cell cultures, acquire a seven-plane z-stack with a z-spacing of 0.2 µm with a 63x or 100x objective, centred on the mitochondrial network. All images within a given experiment must be acquired using identical imaging parameters, including objective lens, laser power, detector gain, exposure time, pinhole size (where applicable), pixel dwell time, and camera settings, pixel size and, where applicable, computational reconstruction setting, to ensure comparability across samples and conditions. Care should be taken to avoid detector saturation in all channels.
2. For 2D heteroplasmy analysis of ΔH2.1 cells (*see* **Section 3.7.2**.), acquire four-channel images using the following channel configuration: Channel 1 - common probe-set signal (CP); Channel 2 - deletion probe-set signal (DP); Channel 3 - Mitochondrial marker (e.g. TOMM20, T20); Channel 4 - Nucleoid marker (e.g. TFAM, TF). Sequential channel acquisition is recommended to minimise spectral cross-talk between fluorophores. Because mtDNA-smFISH puncta are at or near the diffraction limit, images should be sampled sufficiently finely to resolve individual puncta without introducing sampling-dependent merging. For conventional diffraction-limited fluorescence imaging using high-NA 63×–100× objectives, a specimen-plane pixel size of approximately 50–100 nm provides a practical starting range, depending on the objective numerical aperture, emission wavelength and microscope configuration. Users should determine the appropriate sampling for their own optical configuration rather than reproduce the nominal pixel dimensions used here. For the imaging configurations described in this protocol, z-stacks were acquired with an axial spacing of 0.2 µm. When computationally reconstructed imaging approaches such as Airyscan are used, the reconstruction procedure should be considered part of the image-processing workflow. Reconstruction settings can influence the apparent intensity, size, contrast and separation of punctate objects and should therefore be kept constant across samples and experimental conditions. Spot-detection and other quantitative analysis parameters should be validated for the reconstructed image type rather than transferred directly from conventional confocal or spinning-disk datasets.
3. To image *Drosophila* tissues, acquire several stacks and ensure step size is no greater than 0.3 µm and that the slice thickness is no greater than 1µm, to ensure proper reconstruction.

### 3.7 Image analysis

#### 3.7.1 3D mtDNA-smFISH analysis

1. Create a 3D cell mask in Fiji ^38^ by cropping stacks to single cells
2. Use MorphoLibJ ^39^ to segment based on a membrane marker. Alternatively, segmentation can be done manually or through alternative segmentation platforms (*see* **Note 21**).
3. Quantify mtDNA-smFISH puncta with FISH-quant ^40^ using cell masks as input
4. For cells which reliable segmentation is difficult to obtain, mtDNA-smFISH puncta can be counted manually in Fiji using CellCounter. ^38^

#### 3.7.2 2D mtDNA-smFISH analysis

To estimate heteroplasmy levels of single-cells or regions of interest, image processing and analysis can be performed in 2D, on single sections or maximum intensity projections, in FiJi using custom macros (**Figure 5**). General considerations for reproducible image analysis are provided (*see* **Note 22**). A companion analysis worksheet summarising recommended parameters, common analytical pitfalls, and good practice guidelines is provided in **Table 2**. All 2D mtDNA-smFISH analysis scripts are freely available (*see* **Note 23**).

1. Generate maximum intensity projection of each z-stack to consolidate near-diffraction-limited nucleoid puncta across planes, improve signal-to-noise ratio, and avoid slice-to-slice redundancy during 2D spot detection and colocalization. For datasets in which axial separation of puncta is important, retain and analyse the original z-stack using the 3D workflow (Section 3.7.1; see Note 22). Go to: [Image → Stacks → Z Project. Select projection type: Maximum Intensity].
2. Isolate regions of interest (ROIs) manually as individual cells using polygonal selection tools. To clear pixels outside the ROI, go to: [Edit → Clear Outside].
3. Channels for probe signal are processed equally to enhance single nucleoid visualisation by removing background fluorescence using a rolling ball algorithm and an initial Gaussian blur. Go to: [Process → Subtract Background (rolling ball radius = 50 pixels). Process → Filters →Gaussian Blur (σ = 2)].
4. To better discern single mtDNA-smFISH foci, a Laplacian-based enhancement filter is applied by using a 5×5 kernel with centre weight of 24 and surrounding weight of -1, through convolution filtering with normalisation. Go to: [Process → Convolve → Laplacian (centre = 24; surrounding = -1)].
5. To remove spurious peaks, a morphological minimum filtering is applied, followed by a final Gaussian blur for smoothing. Go to: [Process → Filters → Minimum (radius = 2 pixels). Process → Filter → Gaussian Blur (σ = 2)].
6. Process the mitochondrial marker (T20) channel (e.g. anti-TOMM20), to remove background before using it as a mask to restrict the analysis to mitochondrial regions. To do so, use a rolling ball algorithm and Gaussian blur. Go to: [Process → Subtract Background (rolling ball radius = 50 pixels). Process → Enhance Contrast (saturated pixels = 0.35%). Process → Filters → Gaussian Blur (σ = 2)].
7. Binary masks using the T20 channel are then created using Otsu’s automatic thresholding. Go to: [Image → Adjust → Threshold (Otsu). Apply → Create Binary Mask].
8. (Optional) Steps 6 and 7 can also be applied to the nucleoid marker channel (TF) if required, e.g. using an anti-TFAM immunostaining.
9. Processed probe channels are masked with the mitochondrial marker (T20) mask created in Step 7, creating two parallel versions: intensity-preserved 32-bit images for quantitative analysis and user-adjustable thresholded versions for visualisation. Probe-positive puncta located outside the T20-defined mitochondrial region are excluded from subsequent spot detection and quantification, irrespective of their fluorescence intensity. Go to: [Process → ImageCalculator → Multiply]
10. Perform spot detection using the Find Maxima algorithm. Go to: [Process → Find Maxima (CP prominence = 15; DP prominence = 12)]. Colocalised spots can be identified through coordinate-based proximity analysis with a three pixel (300 nm) limit.
11. For estimation of heteroplasmy levels in cells carrying a single mtDNA deletion (as in the ΔH2.1 cells), this is inferred as:

**Figure 5.**
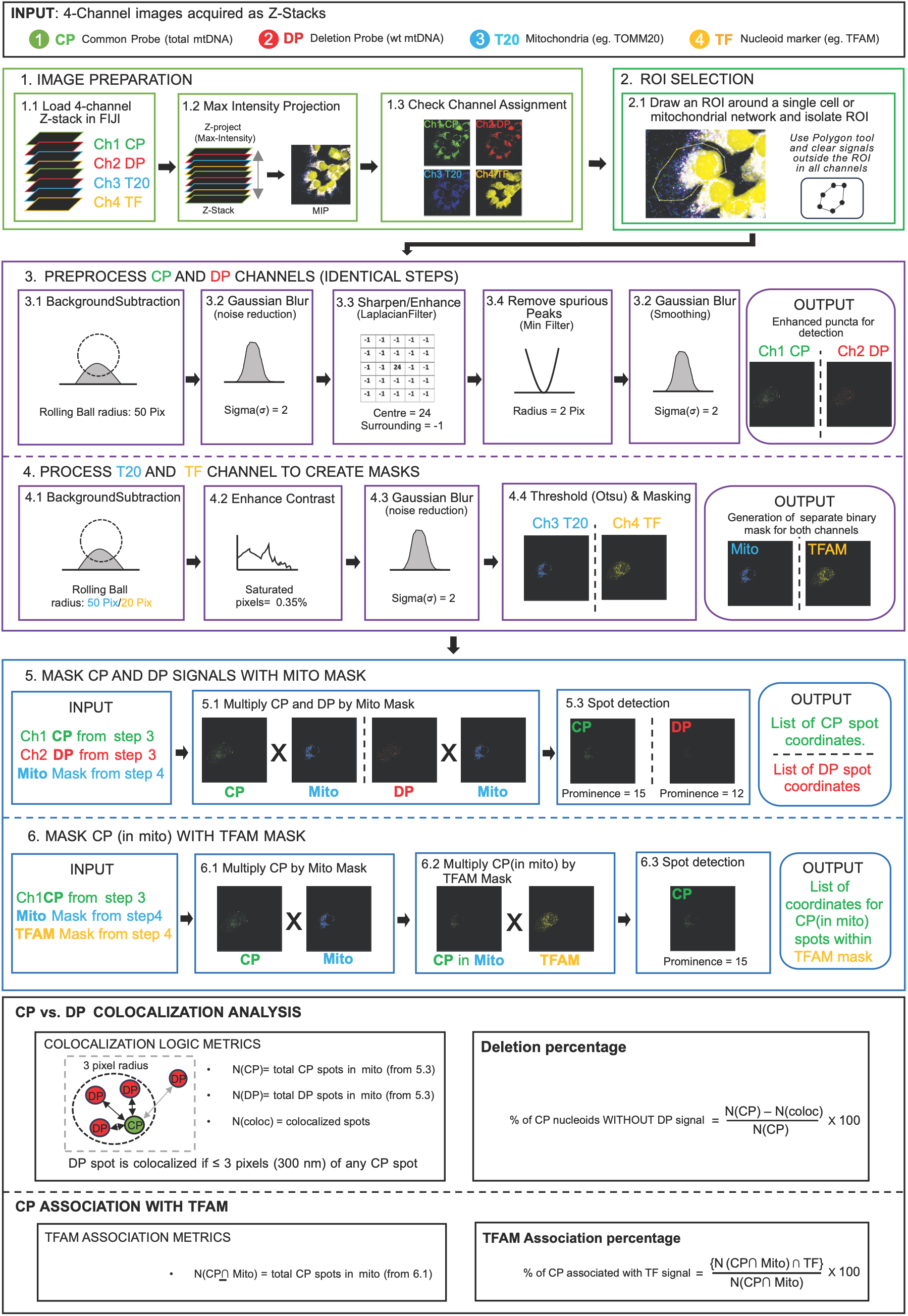
2D mtDNA-smFISH Image analysis pipeline. Workflow for quantification of mtDNA-smFISH signals from 2D fluorescence microscopy datasets, as detailed in **Section 3.7.2**. Four-channel image stacks containing common probe (CP), deletion probe (DP), mitochondrial marker (T20; e.g. TOMM20), and nucleoid marker (TF; e.g. TFAM) signals are processed in Fiji using custom analysis scripts. Images are analysed either as single optical sections or, typically, as maximum intensity projections (MIPs) (*see* **Note 22**). Following manual selection of regions of interest (ROIs), CP and DP channels undergo background subtraction, smoothing, Laplacian-based enhancement, and spot refinement to improve detection of individual mtDNA-smFISH puncta. Mitochondrial and nucleoid marker channels are processed independently to generate binary masks that restrict analysis to biologically relevant regions. Probe signals are subsequently masked and subjected to coordinate-based spot detection and colocalization analysis. Heteroplasmy is estimated from the proportion of CP signals lacking associated DP signal, while association of mtDNA-smFISH puncta with nucleoid markers is quantified through sequential mask application and spot counting.

**Table 2.** Companion worksheet for 2D mtDNA-smFISH image analysis. . Accompanying **Figure 5** and **Section 3.7.2** describing recommended image processing parameters, common sources of analytical variability, and practical guidance for reproducible implementation of the mtDNA-smFISH analysis workflow, intended as an operational reference and quality control aid, to be used alongside the protocol and image analysis scripts. Spatial conversions are based on the confirmed specimen-plane pixel size of 100 nm/pixel for the 2D analysis datasets. Values should be treated as starting settings for comparable data and validated when transferred to other optical configurations or image types. Note: The 500 nm value for the Laplacian filter describes the specimen-plane footprint of the 5 × 5 pixel kernel. It should not be interpreted as a resolution limit or cutoff frequency.

| **Processing step** | **Parameter** | **Value used** | **Physical scale** | **Adaptation guidance** |
|---|---|---|---|---|
| Background subtraction (CP/DP) | Rolling-ball radius | 50 px | 5 µm | Preserve the approximate physical radius when resampling; validate background removal without suppressing puncta. |
| Background subtraction (TF) | Rolling-ball radius | 20 px | 2 µm | Use as a starting value; validate for TF signal morphology and background characteristics. |
| Gaussian blur | Sigma (σ) | 2 px | 200 nm | Rescale with specimen-plane sampling where appropriate; confirm that smoothing does not merge adjacent puncta. |
| Laplacian enhancement | Kernel footprint | 5 × 5 px; centre = 24, surrounding = -1 | 500 × 500 nm footprint | Kernel weights are dimensionless. The physical footprint may be adapted to sampling, then validated against punctum morphology. |
| Minimum filter | Radius | 2 px | 200 nm | Rescale with sampling as a starting point and verify that true puncta are retained. |
| Threshold generation | Thresholding algorithm | Otsu | N/A | Algorithmic choice rather than a spatial parameter; validate mask quality for each image type. |
| Find Maxima (CP) | Prominence | 15 | N/A | Intensity-dependent; re-optimise empirically if acquisition, reconstruction, labelling or intensity scaling changes. |
| Find Maxima (DP) | Prominence | 12 | N/A | Intensity-dependent; re-optimise empirically if acquisition, reconstruction, labelling or intensity scaling changes. |
| CP-DP colocalisation | Distance threshold | ≤3 px | ≤300 nm | Use the physical distance as a starting criterion and validate against optical resolution, registration and centroid uncertainty. |

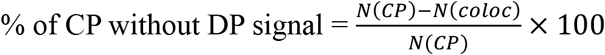

12. To calculate the association with a nucleoid marker, apply the nucleoid marker channel mask (see Step 8), against the mtDNA-smFISH signal and calculate:

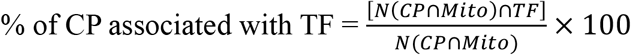

## 4 Notes

1. Virtually equivalent disposables, reagents, instruments and software can be purchased from multiple other providers at a similar cost.
2. Alternatively, solutions can be prepared as follows: ^26^
  - Probe hybridisation buffer: 30% formamide, 5x sodium chloride sodium citrate (SSC), 9 mM citric acid (pH 6.0), 0.1% Tween-20, 50 mg/mL heparin, 1x Denhardt’s solution, 10% dextran sulfate
  - Probe wash buffer: 30% formamide, 5x sodium chloride sodium citrate (SSC), 9 mM citric acid (pH 6.0), 0.1% Tween-20, 50 mg/mL heparin
  - Amplification buffer: 5x sodium chloride sodium citrate (SSC), 0.1% Tween-20, 10% dextran sulfate
3. Sequences of probe halves can be ordered individually from any commercial provider of oligonucleotides. We tend to request HPLC purification when ordering probes, to ensure correct sequence composition, and increase assay specificity, but have not formally compared how different oligonucleotide purification methods affect mtDNA-smFISH signal specificity. As oligonucleotides come, or are usually resuspended, at 100 µM, it is recommended to prepare a working solution at 1 µM for easier dilution to reach the final concentration in the protocol (typically 4 nM).
4. The target sequence reverse complement can be easily inferred at http://reverse-complement.com.
5. Probes were designed with initiator sequences compatible for different amplifier hairpins provided by Molecular Instruments (**Table 3**) (all 5’-3’). It should be noted that the most recent chemistry and sequences used by Molecular Instruments (‘Gold’) remain proprietary, and have as yet not been tested as part of this protocol.
6. Probe hybridisation efficiency to its target sequence is likely to depend on various factors, such as the number of probes, probe length, GC content, intrinsic secondary structures, and proteins bound to the target sequence *in situ*. Target sequences can be determined together with a commercial provider (see **Reagents**). Alternatively, the Stellaris probe designer (https://oligos.biosearchtech.com/products/rna-fish/stellaris-probe-designer) or an open-source smFISH-HCR probe designer software ^36^ have been used successfully to identify appropriate target sequences. The more probes used, the stronger the signal. We typically use 10 probe pairs for mtDNA-smFISH, but good results can be obtained with as little as three probes, and even with a single probe-pair (*see* **Note 15**). We aim to have a GC content around 40-60%, but have obtained good results with GC contents as low as 15%. We have so far not tried probe lengths other than 25 nucleotides.
7. To choose target sequences that can distinguish different heteroplasmic mtDNA molecules within the same cell, we have mostly relied on small or large indels. The smallest sequence divergence we have tested so far contains 6 mismatches in one probe-half. We usually try to include as much sequence divergence as possible within one half of the probe pair, rather than distributing the divergent nucleotides across both halves. We have so far not yet tested whether mtDNA-smFISH can distinguish single-nucleotide polymorphisms.
8. To ensure the absence of non-specific signal coming from probe recognition of mRNAs, RNase treatment can be performed following the de/rehydration steps (*see* **Section 3.3.2**). ^36,41^ However, the harsh denaturation conditions of mtDNA-smFISH are likely to result in degradation of most RNA. Specificity of the signal (*see* **Notes 9, 10**) can further be tested by DNAse treatment of the samples at this stage as well, as previously described by others ^36^.
9. Specificity of the probes for mtDNA in cell lines can be tested in different ways. We typically generate Rho^0^ cells, derived through 2’-3’-dideoxycytidine (ddC) (Sigma-Aldrich D5782) treatment, resulting in mtDNA depletion. ^42,43^ Cells must be treated with 500 µM ddC for two weeks with frequent media changes. Alternatively, mtDNA-smFISH can be conducted on cells of a different species with a different mtDNA sequence (e.g. human probes on mouse or *Drosophila* cells, and vice versa). DNase treatment of the cells prior to mtDNA-smFISH is an additional option to determine specificity of signal for DNA, and can be conducted in parallel to the mtDNA-smFISH on the cells of choice. ^36^
10. Specificity of the probes for mtDNA in whole-mount *Drosophila* tissue can be tested by conducting mtDNA-smFISH on tissue of different *Drosophila* species with a different mtDNA sequence (e.g. *D. melanogaster* probes on *D. yakuba* tissue, and vice versa). Alternatively, mtDNA depletion in cells within a tissue can be achieved genetically, through GAL4-UAS-mediated expression ^44^ of a mitochondrial-targeted restriction enzyme recognising the mtDNA sequence. For example, expression of mito-XhoI (*see* **Section 2.2.4**) ^33^ in *Drosophila* NSCs using a NSC-specific GAL4-line (*see* **Section 2.2.4**) depletes WT *D. melanogaster* and *D. yakuba* mtDNA. ^29,45^
11. We have consistently observed low-level non-specific signal with the *D. yakuba*-specific probes. We have not been able to identify the exact reason, but presume this is due to cross-reactivity with *D. melanogaster* mtDNA. ^29^ In our experience, this can be addressed by adjusting thresholds for signal quantification, based on the values obtained with the *D. yakuba* probe set in *D. melanogaster* tissue (*see* **Note 10**).
12. For straightforward genotyping of heteroplasmic *Drosophila* strains carrying both *D. melanogaster* and *D. yakuba* mtDNA, we perform PCR on DNA extracted from whole flies using the following primers: forward primer: 5’-ttagaccaaatttattgggagaccc-3’; reverse primer: 5’-ggaattcctcaacctttttgtgatgc-3’. The target is a region of mtDNA shared between *D. melanogaster* and *D. yakuba* that differs in the presence of an NdeI restriction site in *D. melanogaster*, but not in *D. yakuba*. The amplicon is approximately 1.25 kb, corresponding to m.11228-12511 on the *D. melanogaster* mtDNA. We digest the resulting PCR product with NdeI, which is active in most PCR buffers (*see* https://www.neb.com/en-gb/tools-and-resources/usage-guidelines/activity-of-restriction-enzymes-in-pcr-buffers) and run it on an agarose gel for visualisation. NdeI will only cut *D. melanogaster* mtDNA, and should therefore produce two bands of 800 bp and 450 bp, while the amplicon from *D. yakuba* mtDNA will remain as a single band of 1.25 kb. Band intensities can be compared in Fiji, or on an automated fragment analyzer for heteroplasmy estimation.
13. The optimal number of cells to seed should be adjusted depending on the cell line in use and accounting variability in culture handling between operators. Aim for a 70-80% confluency at the time of fixation.
14. Lower volumes of reagents can be used, provided that proper coverage of the sample is ensured and evaporation is avoided.
15. It is recommended to keep excess tissue attached to the tissue of interest, to make sure the tissue remains visible in the vials. In addition, the frequent washes in the subsequent steps may lead to tissue loss, and it is recommended to dissect more samples than required for the downstream analysis.
16. Many antibodies require target antigens to be in their native folded state for binding and detection. As mtDNA-smFISH requires heat denaturation of the samples, native protein structure is disrupted, and some antibodies will no longer work for immunostaining. To detect epitopes that do not withstand heat denaturation, proceed with normal primary antibody staining, then use a biotin-conjugated secondary antibody, and fluorophore-conjugated streptavidin. Antibody steps must be performed prior to the denaturation step (*see* **Section 3.3.3**) of mtDNA-smFISH, and streptavidin must be added following the amplification and serial washing steps (*see* **Section 3.4.2**). Streptavidin will bind to the biotin on the secondary antibody allowing fluorescent detection of the desired target, as described previously. ^29,37^ If high background signal from endogenous biotin is an issue, commercially available streptavidin/biotin blocking kits may help.
17. Several of the antibodies we used continue to bind antigens in cell culture or whole-mount tissue after denaturation. These include the antibodies listed in the Reagents (*see* **Section 2.6**), against ATPsynα, FLAG, TOMM20 and TFAM. However, immunostaining quality may be affected, with decreased signal-to-noise ratio or structural changes due to destruction of the cellular ultrastructure. ^37^ Alternatively, antibodies have also been raised specifically to heat-denatured antigens, such as for GFP. ^35^
18. Probes can be tested individually to ensure their specificity. To do so, a higher concentration is advised. As an example, individual testing of the probe sets reported here was previously performed at 400 nM. ^29^
19. The time required for HCR amplification will vary and needs to be optimised for probes and conditions. We typically allow a 6-hour amplification time for cells in culture with at least 10 probes, but increase this, for example to 12 hours, for whole-mount tissue or when there are fewer probes.
20. Any mounting medium can be used, in line with routine laboratory practices. We typically mount cultured cells with multiple drops of DAKO fluorescent mounting medium, with a refractive index of 1.372, and whole-mount tissues in Prolong Diamond antifade mounting medium, with a refractive index of 1.47 upon curing. When using a curing mounting medium, ensure that it is fully cured for a minimum of 24 hours in the dark before sealing. Some curing mounting media may lead to tissue shrinkage.
21. Several tools are available for automated 3D segmentation and single-cell assignment in tissues, and can be integrated with FISH-quant v2 ^40^ to associate cell identities with mtDNA-smFISH puncta. For example, Cellpose ^46^ is a freely available deep-learning-based cell segmentation framework implemented in Python that enables 3D segmentation using nuclear, membrane, or other markers (i.e. DAPI or membrane-targeted GFP) that are compatible with mtDNA-smFISH. Segmentation outcupts can be incorporated into FISH-quant v2 (big-fish) analysis workflows to enable automated 3D quantification of heteroplasmy at single-cell level in complex tissue volumes. More broadly, coupling automated segmentation with mtDNA-smFISH analysis frameworks may enable future cell-resolved and spatially contextualised heteroplasmy mapping across intact tis-sue volumes, supporting scalable analysis of mitochondrial genetic diversity at tissue and population scales.
22. For reproducible quantification of heteroplasmy and colocalization using 2D mtDNA-smFISH analysis, image processing parameters should be applied consistently across samples and experimental conditions. Analysis is generally performed on maximum intensity projections (MIPs) unless individual optical sections are specifically required, as projection of z-stacks consolidates near-diffraction-limited nucleoid signals across planes, improves signal-to-noise ratio, and reduces z-registration artefacts during spot detection and colocalization measurements. MIP also removes axial information and can cause puncta located at different positions along the z-axis to appear overlapping or merged in the projected image. Consequently, closely spaced puncta may be detected as a single object, and signals from different channels that are separated axially may appear colocalised after projection. For samples with substantial axial depth, high puncta density, or where accurate three-dimensional spatial association is required, analysis of the original z-stack using the 3D workflow described in Section 3.7.1 is recommended. Quantification should be performed on individual cells or manually defined regions of interest (ROIs) rather than full fields of view. Probe-positive puncta located outside the T20-defined mitochondrial mask are excluded from quantitative analysis. Such extra-mitochondrial puncta should not be classified as mtDNA nucleoids on the basis of fluorescence intensity alone, irrespective of their apparent signal intensity. Quantitative analysis should be carried out using intensity-preserved 32-bit images (or 16-bit where appropriate), while thresholded representations should be reserved for visualisation and mask generation only.
23. 2D mtDNA-smFISH analysis scripts are available on GitHub (https://github.com/JvdAlab/mtDNA-smFISH/tree/main/imaging_analysis).

## 5 Concluding remarks

Accurate sequence-specific visualisation of mtDNA molecules *in situ*, within cells and tissues, is a prerequisite for the in-depth investigation of mtDNA dynamics during development and ageing. The mtDNA-smFISH protocol presented here provides a robust method for the visualisation of mtDNA variants using single-molecule FISH combined with enzyme-free HCR-mediated signal amplification. We demonstrated the utility of mtDNA-smFISH in cell culture, using different cell lines from different species, and in whole-mount *Drosophila* tissues. A versatile co-immunostaining protocol, compatible with the harsh denaturation conditions, opens the possibility for future studies of genetic mtDNA variation at subcellular or single-mitochondrion resolution.

**Table 3:**
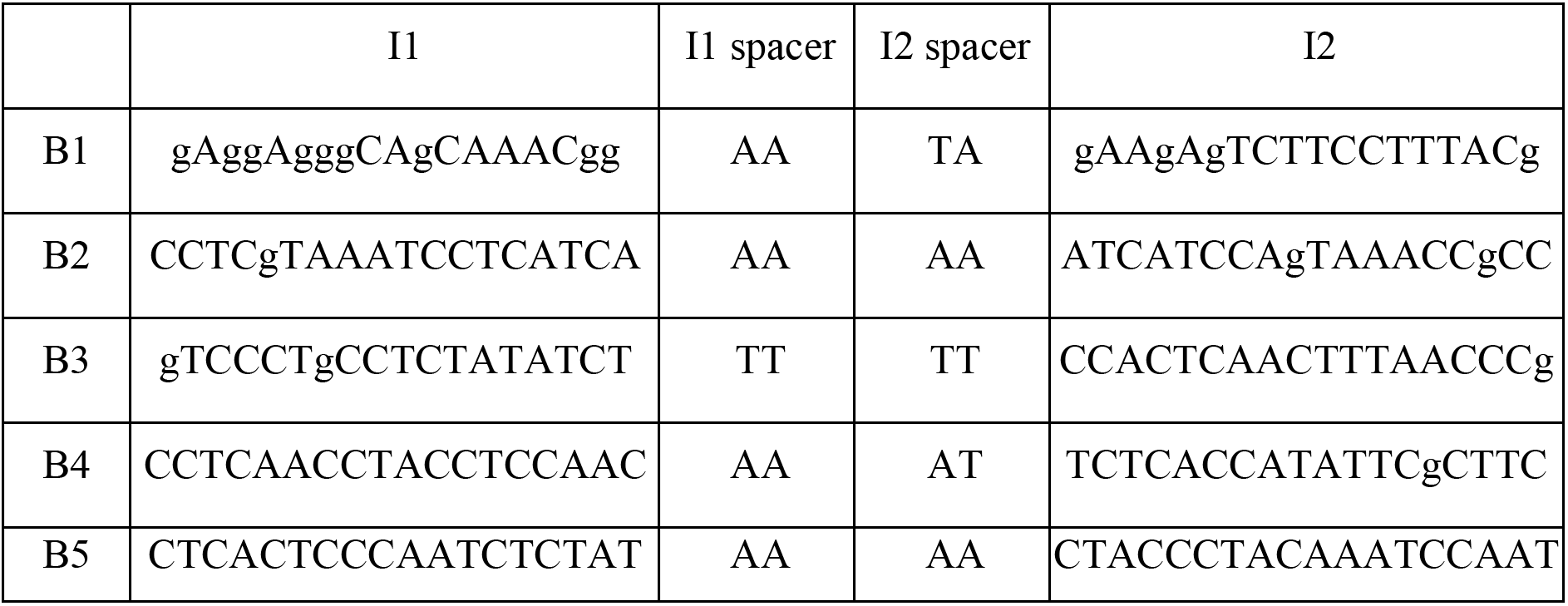
Initiator sequences compatible for different amplifier hairpins provided by Molecular Instruments. All sequences provided are 5’-3’. (*see* Note 5)

To ensure efficient and accessible mtDNA-smFISH image analysis, we developed a pipeline in Fiji for 2D heteroplasmy quantification. While existing software for 3D smFISH-analysis allows accurate heteroplasmy quantification by better accounting for the full spatial distribution of mtDNA signals, most packages are either not freely available or widely and readily accessible to the broad research community. Our 2D heteroplasmy pipeline instead serves as a straightforward tool for estimating mutation burden and comparing this between cells, but has the advantage of relying exclusively on open-access software, making it more accessible and easier to implement.

In order for mtDNA-smFISH to distinguish different mtDNA molecules and maximise probe specificity, we have relied on the presence of large-scale deletions, or several single-nucleotide differences or indels. However, precisely how much sequence divergence is required remains to be determined, and we have so far not tried to differentiate mitochondrial genomes that differ only by a single-nucleotide polymorphism. Further improvements and modifications to this protocol may help to increase sensitivity and specificity at various stages of the mtDNA-smFISH protocol. For example, through addition of random oligonucleotides during the pre-hybridisation and hybridisation steps. ^47^ A major future goal will then be to develop single-nucleotide mtDNA-smFISH approaches capable of detecting defined point mutations, thereby expanding the applicability of this method to a broader range of pathogenic and naturally occurring mtDNA variants.

Beyond the quantification of heteroplasmy, mtDNA-smFISH should provide new opportunities to investigate the pathways that control mtDNA cell biology more broadly, including mechanisms regulating mtDNA maintenance, degradation, intracellular distribution and trafficking, and release into the cytosol to trigger inflammatory signalling. ^48–52^ This will be particularly valuable for understanding the distinct behaviours and fates of mtDNA molecules within cells and tissues. Indeed, individual mtDNA molecules may exert local effects, or “spheres of influence”, within the mitochondrial network and surrounding cellular environment. ^53^ This process could drive inner mitochondrial membrane alterations, triggering an inner mitochondrial membrane quality control process and therefore the degradation of selective compartments containing different types of mtDNA. ^54^ In this context, the ability to visualise mutant and wild-type mtDNA in fixed cells already provides a powerful entry point not only to define how WT and mutant mtDNA species behave and localise within cells, but also to determine their local mitochondrial environment.

Because current mtDNA-smFISH approaches are restricted to fixed-cell imaging, they capture static snapshots rather than dynamic behaviours. Novel methodologies will be needed to visualise WT and mutant mtDNA dynamics and trafficking in real time. Such developments would provide important insight into how mtDNA dynamics, quality control and inflammatory signalling are coordinated in living cells. Despite these limitations, mtDNA-smFISH serves as a powerful tool for *in situ* mtDNA analysis and provides a foundation for further development and broader applications, to ultimately unravel the mechanisms that shape mtDNA inheritance and evolution in a wide range of physiological and pathological contexts.

## Acknowledgements

We thank all lab members for helpful discussions and advice; C. Moraes for the ΔH2.1 cybrid cell line; K.C. Nakamura and H. Hioki for anti-heat-denatured-GFP antibodies; H. Ma for *Drosophila* stocks; H. Choi and colleagues from Molecular Instruments for reagents and advice; *Drosophila* stocks obtained from the Bloomington Drosophila Stock Center (NIH P40OD018537) were used in this study.

This work was supported by a Wellcome Clinical Research Career Development Fellowship (219615/Z/19/Z), an Evelyn Trust Medical Research Grant (21–25), a UKRI BBSRC Responsive Mode Research Grant (BB/X00256X/1), a Wellcome Discovery Award (226653/Z/22/Z), a Medical Research Council (MRC) award (MC_PC_21046) to establish a National Mouse Genetics Network Cluster in Mitochondrial Diseases (MitoCluster), the LifeArc Centre for Rare Mitochondrial Diseases (LAC-TreatMito) under grant no. 10748, Rosetrees (PGL23/100048), and MDUK (23SI-PRG60-0013). LifeArc is a charity registered in England and Wales under no. 1015243 and in Scotland under no. SC037861. J.P. and J.v.d.A. acknowledge core funding from the UKRI MRC to the MRC Mitochondrial Biology Unit (MC_UU_00028/5 and MC_UU_00028/8). J.P. was also supported by State funding from the Agence Nationale de la Recherche under France 2030 program (ANR-10-IAHU-01).

For the purpose of open access, the authors have applied a Creative Commons Attribution (CC BY) license to any author-accepted manuscript version arising from this submission.

